# Sensitive colorimetric ELISA measuring salivary SBP-specific secretory IgA, its application to pollinosis SLIT

**DOI:** 10.64898/2025.12.02.691766

**Authors:** Tetsuro Yamamoto, Fusako Mitsunaga, Atsushi Kotani, Kazuki Tajima, Muneki Hotomi, Shin Nakamura

**Affiliations:** Innovation Research Center, EPS Holdings, Inc.; 2-3-19 Kouraku, Bunkyo-ku, Tokyo, 112-0004, Japan; EP Mediate Co., LTD; 6-29 Shin-ogawamachi, Shinjuku-ku, Tokyo, 162-0814, Japan; Research Center, EPS Innovative Medicine Co., Ltd.; 6-29 Shin-ogawamachi, Shinjuku-ku,Tokyo, 162-0814, Japan; Intelligence and Technology Lab Inc., 52-1 Fukue, Kaizu-cho, Kaizu, Gifu, 503-0628, Japan; Biomedical Institute, NPO Primate Agora, 52-2 Fukue, Kaizu-cho, Kaizu, Gifu, 503-0628, Japan; Department of Otorhinolaryngology-Head and Neck Surgery, Wakayama Medical University, Kimiidera 811-1, Wakayama, 641-8509, Japan

**Keywords:** Allergen-specific SIgA, saliva, amplified colorimetric ELISA, pollinosis, sublingual immunotherapy

## Abstract

**Background:** Sublingual immunotherapy is safe and one of the most effective etiological treatments for pollinosis. Salivary allergen-specific secretory immunoglobulin A (SIgA) is a useful marker to objectively evaluate treatment effectiveness. However, saliva samples usually exhibit high background noise because of nonspecific binding in enzyme-linked immunosorbent assays (ELISAs).

**Objective:** We aimed to establish an amplified ELISA to measure allergen-specific salivary SIgA levels to assess the efficacy of sublingual immunotherapy (SLIT).

**Methods:** Biotinyl tyramide (BT-Tyr) was used to amplify the signal in a colorimetric ELISA. The allergen sugi basic protein (SBP) was biotinylated and used as a reporter to obtain sensitive and specific results with low background. This ELISA was used to measure anti-SBP SIgA levels in saliva samples from SLIT-treated patients with Japanese cedar pollinosis.

**Results:** The ELISA was highly sensitive and achieved more than 200-fold signal amplification compared to non-amplified conditions. It was also specific for antigens without background noise. The levels of SBP-specific SIgA in saliva increased in patients 3 months after SLIT initiation.

**Conclusion:** We established a sensitive colorimetric ELISA to measure SBP-specific SIgA in saliva using BT-Tyr-based amplification and biotinylated antigen as a reporter. We used this sensitive ELISA to monitor the levels of SBP-specific SIgA in saliva samples from patients with pollinosis receiving SLIT. Increased salivary SBP-specific SigA levels were observed in these patients.

## Introduction

Allergy to Japanese cedar pollen is a prevalent pollinosis in Japan and patients with allergic symptoms have reduced quality of life (1,2). Immunotherapy is a beneficial treatment for patients with pollinosis. Among immunotherapies, sublingual immunotherapy (SLIT) has advantages over subcutaneous immunotherapy in terms of efficacy, safety, and self-administration, without the need for medical care personnel (3,4). As SLIT is expected to increase allergen-specific SIgA levels in saliva and reduce the allergic response (5,6), anti-allergen SIgA in saliva is a useful biomarker for evaluating treatment efficacy.

In this study, we established a sensitive colorimetric enzyme-linked immunosorbent assay (ELISA) to measure sugi basic protein (SBP)-specific SIgA in saliva samples from patients with Japanese cedar pollinosis treated with SLIT using CEDARCURE (Japanese Cedar Pollen Sublingual Tablets, Torii Pharmaceutical Co. Ltd., Tokyo, Japan) (4). The sensitivity of the ELISA was markedly amplified by the use of biotinyl tyramide (BT-Tyr) (7) and biotinylated SBP (BT-SBP) as an amplifier and detector, respectively. Amplified ELISA reactions were measured using a chromogenic plate reader at an absorbance of 450 nm. We examined the optimal conditions in detail, especially concentrations of hydrogen peroxide (H_2_O_2_) and BT-Tyr.

## Results and Discussion

Figure 1A shows an outline of the ELISA materials, procedures, and conditions. The primary materials used for the amplified ELISA were as follows. The SBP (BioDynamics Laboratory Inc., Tokyo, Japan) was biotinylated using a Biotin Labeling Kit-NH_2_ (Dojindo Laboratories, Kumamoto, Japan) and a centrifugal tube with a molecular weight cut-off of 10kDa (Amicon Ultra, Merck KGaA, Darmstadt, Germany). The concentration of BT-SBP was determined using a DC Protein Assay (Bio-Rad, Hercules, CA, USA).

**Figure. 1.**
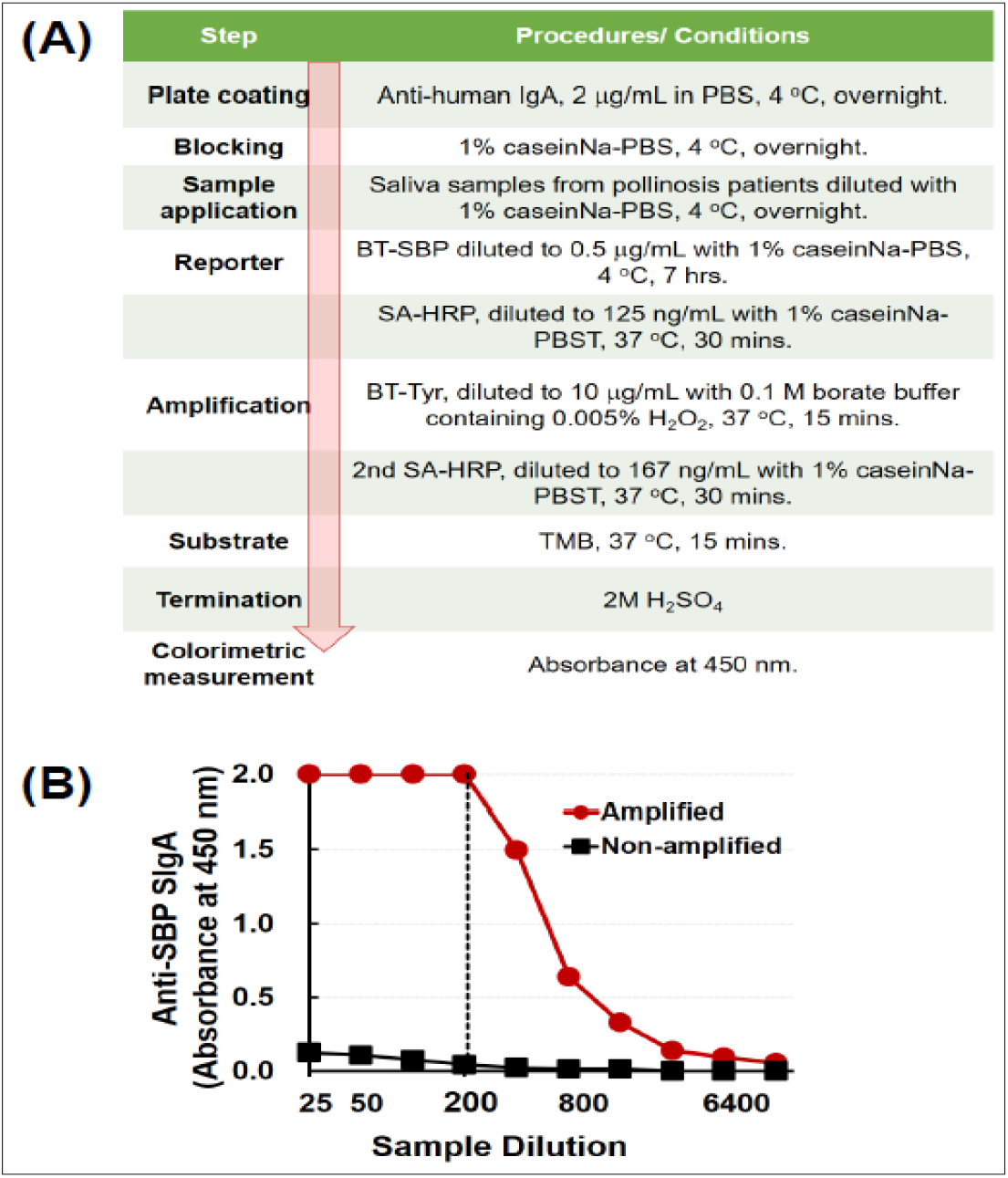
Amplified ELISA for saliva samples utilizing BT-SBP. Outline (A) and sensitivity (B) of the procedure. In the sensitivity test (B), saliva samples from a patient with pollinosis treated with SLIT were serially diluted to a range of 1:25 to 1:12,800. The anti-SBP SIgA in the patient’s diluted samples were measured using the BT-Tyr amplification ELISA (red circles), and compared with non-amplified ELISA (black squares). ELISA, enzyme-linked immunosorbent assay; SBP, sugi basic protein; BT-SBP, biotinylated SBP; BT-Tyr, biotinyl tyramide; SLIT, sublingual immunotherapy; SBP, SIgA, secretory.

BT-Tyr (Sigma-Aldrich, Burlington, MA, USA) was dissolved in dimethyl sulfoxide as a stock solution (50 mg/mL) and divided into several small volumes for one-time use, followed by freezing. The working solution of BT-Tyr was prepared by diluting the stock solution to a final concentration of 100 μg/mL with ethanol, after which, it was kept at 4°C until use. Prior to use of the diluted BT-Tyr, a 30% H_2_O_2_ solution (Fujifilm Wako Pure Chemical Corporation, Osaka, Japan) was diluted to a final concentration of 0.005% with 0.1 M borate buffer (pH 8.5). The diluted BT-Tyr was then added to borate buffer containing H_2_O_2_ to prepare a final concentration of 10 μg/mL, which was used immediately. A BT-Tyr concentration of 10 μg/mL yielded the best amplification, with high sensitivity and low background.

The outline of the procedure and conditions for this amplified ELISA are as follows.

### Plate coating

Polystyrene microplates (Maxisorp, Nunc, Roskilde, Denmark) were coated with anti-human IgA (polyclonal, goat; Mabtech, Stockholm, Sweden) diluted to a final concentration of 2 μg/mL in phosphate-buffered saline (PBS) containing 0.02% sodium azide, pH 7.4. The plates were washed with 0.05% Tween 20-PBS (PBST) at each step, except for the final step which involved the addition of the stop solution to the substrate. Blocking: Plates blocked with casein sodium (casein Na; Fujifilm Wako Pure Chemical Corporation) resulted in a lower background signal than those blocked with bovine serum albumin.

### Sample application

Saliva samples were diluted with 1% casein Na-PBS and then applied to the plates.

### Reporter

BT-SBP diluted to a final concentration of 0.5 μg/mL with 1% casein Na-PBS was added and bound to the immobilized anti-SBP SIgA. Next, streptavidin-conjugated horseradish peroxidase (SA-HRP; Invitrogen-Thermo Fisher Scientific Inc., Waltham, MA, USA) diluted to a final concentration of 125 ng/mL with 1% casein Na-PBST was added.

### Amplification

BT-Tyr diluted to a final concentration of 10 μg/mL with 0.1M borate buffer, pH8.5, containing 0.005% H_2_O_2_ was added, the plates were incubated for 15 min at 37°C, and SA-HRP diluted to a final concentration of 167 ng/mL was added.

### Substrate and terminating reaction

The chromogenic substrate, 3,3′,5,5′-tetramethyl-benzidine (TMB; Sigma-Aldrich) was added to develop the color and the reaction was terminated by adding 2M H_2_SO_4_.

### Colorimetric measurement

The absorbance at 450 nm was measured using a plate reader in the visible light range (iMark Microplate Absorbance Reader, Bio-Rad).

As shown in Figure 1B, the method was highly sensitive and achieved more than 200-fold signal amplification compared to the non-amplified condition. We also confirmed the specificity of the amplified ELISA using an adsorption test, in which several diluted saliva samples were pre-incubated with various concentrations of SBP prior to application to the coated plates.

We used this amplified ELISA to monitor anti-SBP SIgA levels in saliva samples from Japanese patients with cedar pollinosis treated with SLIT. Saliva samples were collected at otolaryngology hospitals from patients with pollinosis treated with or without SLIT at 3-months intervals for 12 months. In the SLIT group, patients were administered 2,000 JAU of CEDARCURE, once daily for the first week after the commencement of treatment. The dose was then increased to 5,000 JAU/d from the second week onward. The patients were carefully examined and observed by doctors during the first administration and in periodic examinations. As shown in Figure 2, the levels of SBP-specific SIgA in the saliva increased in patients while receiving SLIT (Figures 2A, 2B and 2C). Although there were individual differences in the pattern, SBP-specific SIgA levels increased within 3 months in patients with SLIT.

**Figure. 2.**
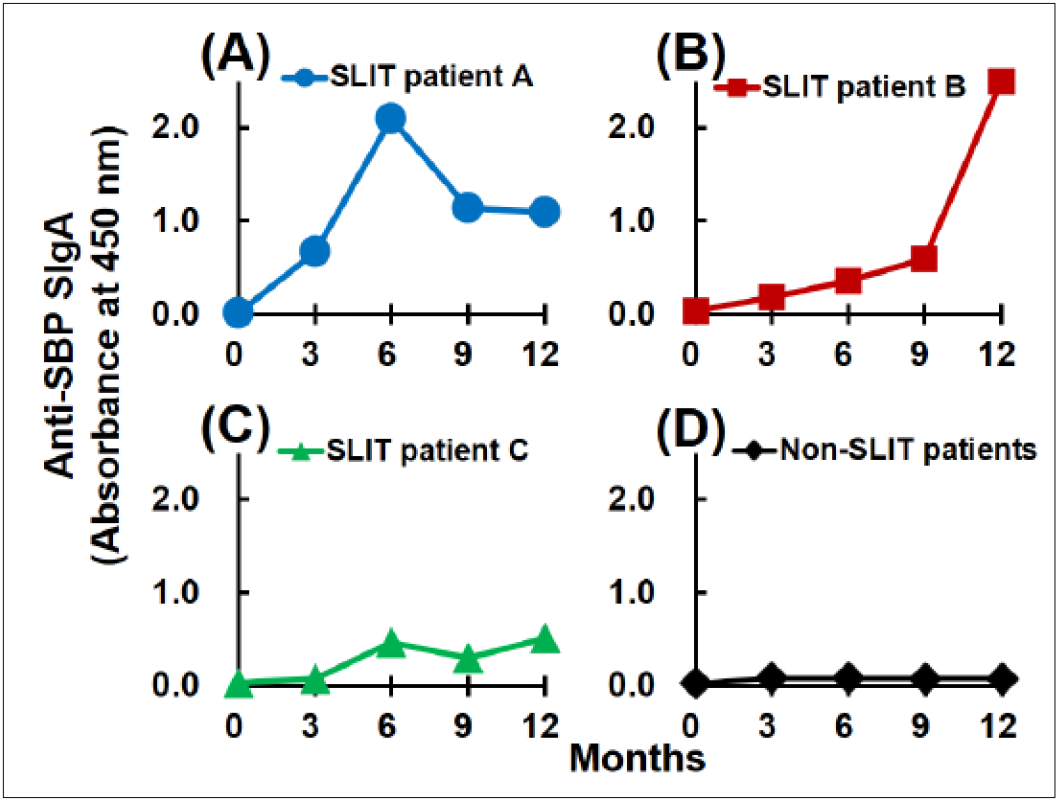
Saliva SBP-specific SIgA in patients with pollinosis followed up over a 12-month period of SLIT. Three different patients with pollinosis who underwent SLIT (A-C). The mean value of three patients with pollinosis treated without SLIT (D).

However, there was no increase in SBP-specific SIgA levels in patients with pollinosis who did not receive SLIT (Figure 2D).

In conclusion, we established a sensitive colorimetric ELISA to measure SBP-specific SIgA levels in saliva using BT-Tyr amplification. We used this sensitive ELISA to monitor the levels of SBP-specific SIgA in saliva samples from patients with pollinosis who received SLIT for 12 months. Using this ELISA, an increase in salivary SBP-specific SIgA levels was observed 3 months after SLIT initiation.

### Institutional Review Board Statement

All sampling procedures were approved by the Research Ethics Committee of Wakayama Medical University (approval number: 3218; date: July 31, 2021). Informed consent and sample collection were conducted under the arrangement of Wakayama Medical University.

## Abbreviations

SLIT: (sublingual immunotherapy),
SB: (sugi basic protein),
BT-Tyr: (biotinyl tyramide),
BT-SBP: (biotinylated SBP).

## Acknowledgement

We thank Kazuhiro Kawai for technical assistance. We also thank Dr. Yuki Shiraishi and Kunihiko Wasaki for their support of this study.

